# Regulated nuclear accumulation of a histone methyltransferase times the onset of heterochromatin formation in *C. elegans* embryos

**DOI:** 10.1101/326231

**Authors:** Beste Mutlu, Huei-Mei Chen, James J. Moresco, Barbara D. Orelo, Bing Yang, John M. Gaspar, Sabine Keppler-Ross, John R. Yates, David H. Hall, Eleanor M. Maine, Susan E. Mango

## Abstract

**ONE SENTENCE SUMMARY:** MET-2/SETDB1 and interactors (LIN-65/ATF7IP and ARLE-14/ARL14EP) initiate heterochromatin formation during embryogenesis.

**ABSTRACT:** Heterochromatin formation during early embryogenesis is timed precisely, but it has been elusive how this process is regulated. Here we report the discovery of a histone methyltransferase complex whose nuclear accumulation determines the onset of heterochromatin formation in *C. elegans* embryos. We find that the inception of heterochromatin generation coincides with the nuclear accumulation of the methyltransferase MET-2 (SETDB). The absence of MET-2 results in delayed and disturbed heterochromatin formation, whereas accelerated nuclear localization of the methyltransferase leads to precocious heterochromatin. We identify two factors that bind to and function with MET-2: LIN-65, which resembles ATF7IP, localizes MET-2 into nuclear hubs, and ARLE-14, orthologous to ARL14EP, promotes stable association of MET-2 with chromatin. These data reveal that nuclear accumulation of MET-2 in conjunction with LIN-65 and ARLE-14 regulates timing of heterochromatin domains during embryogenesis.

## MAIN TEXT

The nucleus of a young embryo undergoes major reprogramming as it transitions from fertilized egg to multicellular embryo. As cells acquire specific fates and zygotic transcription commences, the nucleus is segregated into distinct domains of euchromatin and heterochromatin *(1)*. While much has been learned about the mechanisms that control fate specification and the onset of zygotic transcription *(2, 3)*, little is understood about the processes that establish chromatin domains *de novo* during embryogenesis. To begin to tackle this question, we examined heterochromatin formation in the nematode *C. elegans.*

We began our analysis with a survey of wild-type embryos using two assays for heterochromatin: First, we used transmission electron microscopy (TEM), where heterochromatin domains can be detected as electron-dense regions (EDRs) within nuclei *(4, 5)*. Second, we surveyed histone modifications to track their abundance and morphology in early embryos. By TEM, nuclei at the earliest stages appeared relatively homogenous, with light speckling in the nucleoplasm and a nuclear envelope free of electron-dense material (**Fig. 1A-B**). With the initiation of gastrulation (~21-50 cell stage), embryos gained more electron-dense puncta throughout their nuclei. By mid-gastrulation (51-100 cell stage), dark material was observed abutting the nuclear envelope, and the nucleoplasmic puncta coalesced into larger, but fewer, electron-dense compartments. These features became more pronounced over time, with large EDRs that spanned the nucleus and bordered the nuclear periphery (**Fig. 1B**, >200 cells). We note that EDRs appeared throughout the embryo, suggesting that cells destined to produce different cell types nevertheless generated heterochromatin at about the same time in development (**fig. S1A**).

**Fig. 1.**
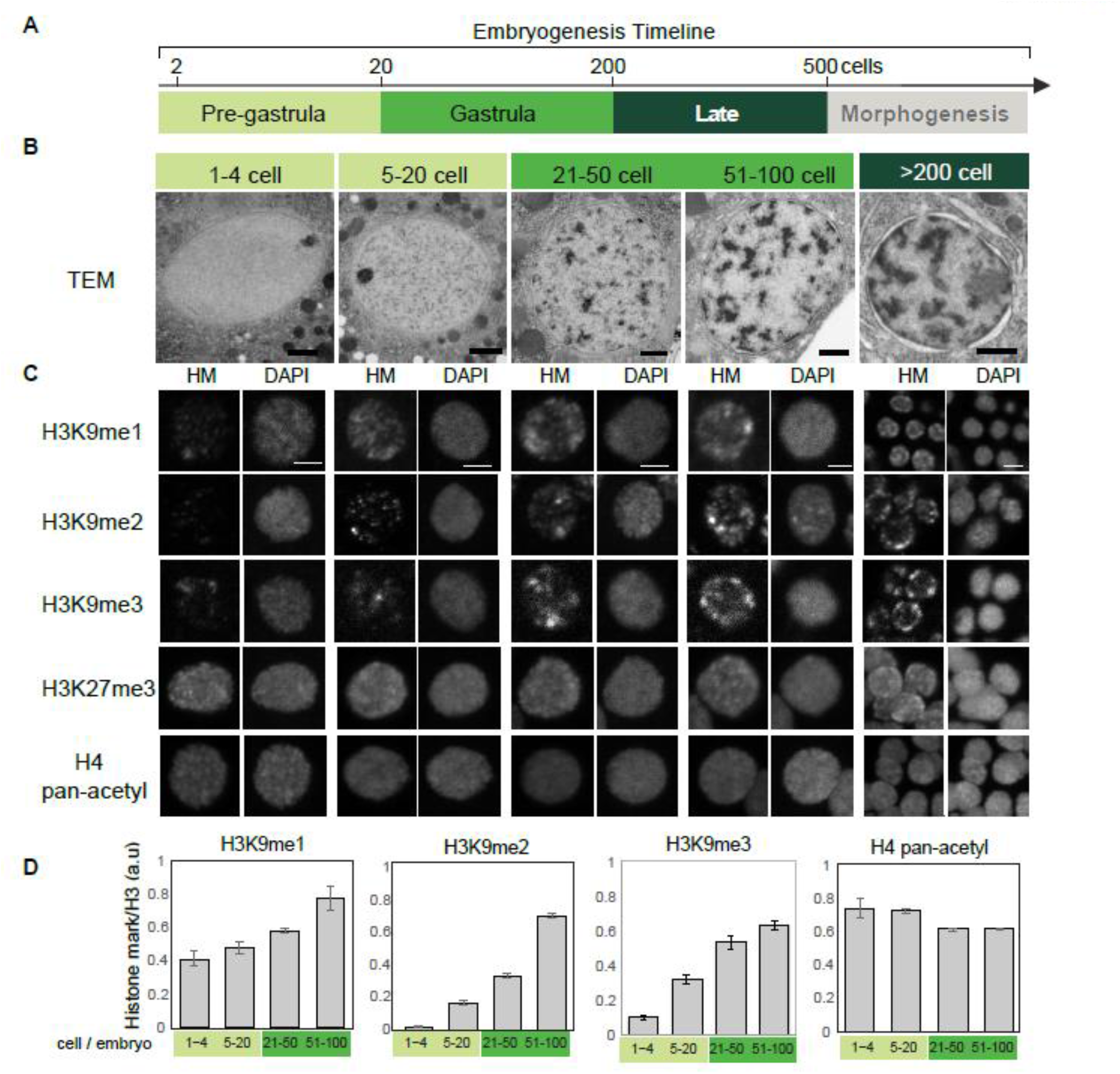
Heterochromatin and H3K9me domains are established during embryogenesis. **A**. Time line of *C. elegans* embryogenesis. Stages are color-coded: light green (<20 cell, pre-gastrula), green (21-200 cell, gastrula), dark green (200-500 cell, late stage). Morphogenesis starts after the 500-cell stage and was not analyzed in this study. **B**. Transmission electron micrographs (TEM) of representative nuclei from wild-type embryos (Scale bar, 1 μm). **C**. Survey of histone modifications. Representative single nuclei at designated embryonic stages stained for histones and DNA (Scale bar, 2 μm). **D**. Quantitation of histone modifications normalized to total histone H3.

Antibody staining revealed a dramatic increase in Histone H3 Lysine 9 methylation (H3K9me) from fertilization to the mid-gastrula during interphase. H3K9me2 increased ten-fold while H3K9me1 and H3K9me3 increased two-fold and six-fold (**Fig. 1C, D**). H3K9me2 was barely detectable at fertilization, but bright puncta became apparent by the 20-cell stage throughout nuclei. The signal intensified during gastrulation with more puncta and brighter staining within puncta (~51 −100-cell stage, mid-stage). A time series of whole embryo stains indicated that most interphase nuclei behaved similarly (**fig. S1B**), and two different antibodies against H3K9me2 gave identical results (**fig. S1C**). The timing and morphology of the H3K9me2/me3 puncta suggest that these modifications are a useful proxy to visualize heterochromatin domains. Both puncta of histone modifications and EDRs arise during gastrulation, beginning around the 20-cell stage. We note that not all histone modifications changed in early embryos: neither levels nor morphology of H3K27me3 or pan-H4Ac were altered (**Figure 1C, D**; see also *(6)*).

To identify the molecular basis of heterochromatin and H3K9me formation, we focused on the methyltransferase MET-2. MET-2 is homologous to vertebrate SETDB1 *(7, 8)*, and required for virtually all H3K9me1 and H3K9me2, as well as some H3K9me3 *(9, 10)*. H3K9me2 was regulated dynamically during embryogenesis, similar to EDRs and its location in the genome, by ChIP, tracks well with heterochromatin proteins such as HPL-2/HP1 *(11)*. Thus, we asked whether loss of MET-2 impacted heterochromatin domains visible by TEM. As a control for TEM fixation and sectioning, we examined cytoplasmic organelles and yolk droplets from *met-2* mutants, which resembled those of wild-type embryos (**fig S2A, B**). Pre-gastrula *met-2* embryos matched their wild-type counterparts, with homogeneous, translucent nuclei (**Fig. 2A**). However, the speckles observed in wild-type nuclei at the 20-cell stage were dimmer in *met-2* mutants, and they failed to coalesce into EDRs by the 50-100-cell stage (**Fig. 2A**). EDRs emerged in older embryos, but they occupied less nuclear volume and were reproducibly paler (**Fig. 2A**). We performed line-scan analysis to quantify the appearance of nuclear EDRs. The standard deviation of line-scan values is higher in nuclei with EDRs compared to homogeneous nuclei that lack heterochromatin. *met-2* nuclei had a homogeneous distribution of signal and a smaller standard deviation at every stage (**Fig. 2B**). These data show that the formation of heterochromatin was disturbed and delayed in *met-2* mutants. These results indicate that *met-2* is critical for the timely formation of segregated heterochromatin domains.

**Fig. 2.**
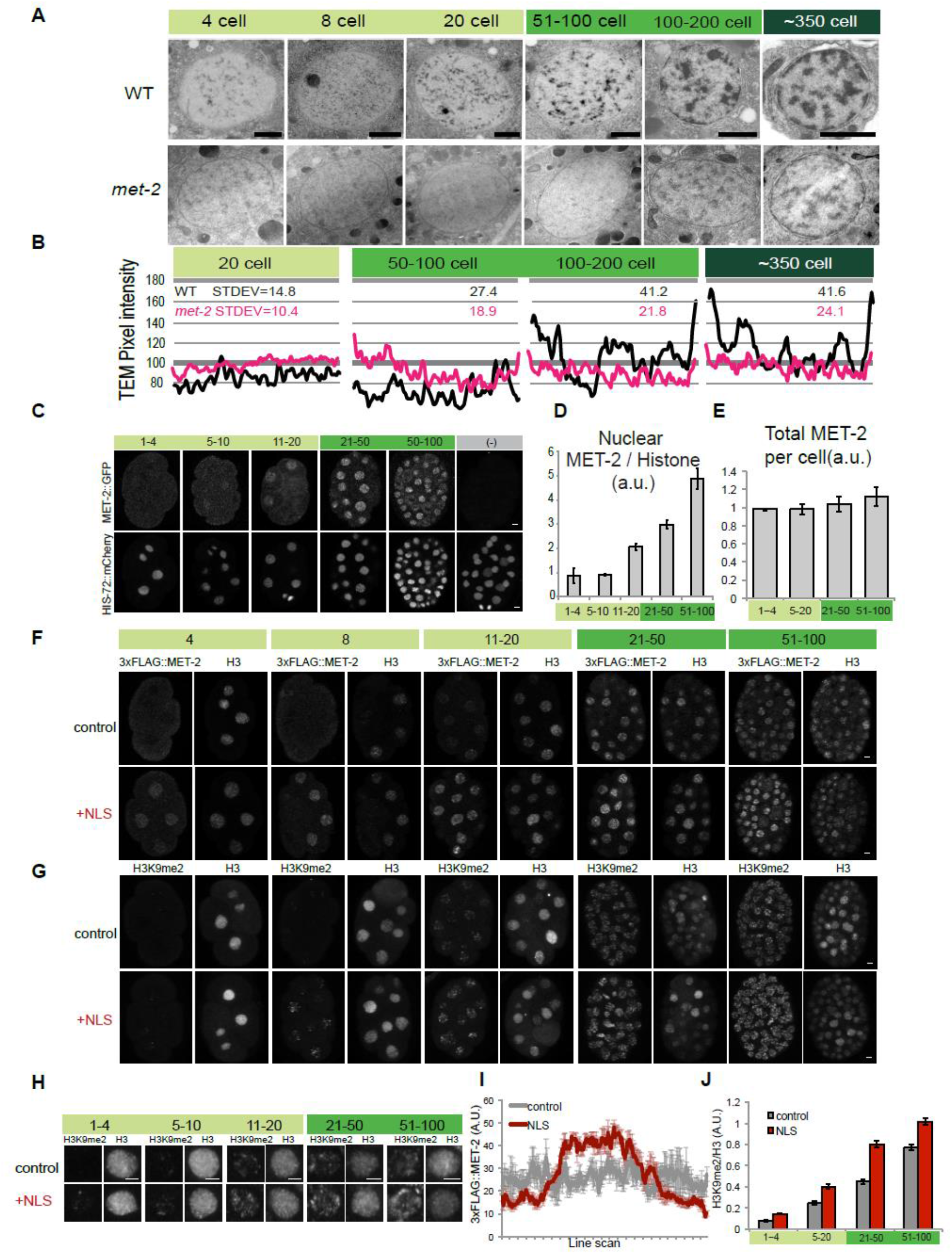
Nuclear accumulation of H3K9 methyltransferase MET-2/SETDB1 determines the onset of heterochromatin formation. **A**. TEM of representative nuclei from wild-type (WT) or *met-2* embryos (Scale bar, 1 μm). **B**. TEM line-scan analysis for WT (black) or *met-2* (magenta) nuclei and the standard deviation (stdev). Line scan for a single nucleus is shown as an example. **C**. Embryos stained for MET-2::GFP (upper) and his-72::mCherry (lower) at designated stages of embryogenesis (Scale bar, 2 μm). **D-E**. Quantitation of nuclear (D) and total (E) MET-2 at designated stages, normalized to his-72::mCherry. **F-G**. Localization of 3xFLAG::MET-2 with a c-Myc NLS compared to 3xFLAG::MET-2 control (F) and corresponding H3K9me2 levels (G). **H**. Interphase nuclei showing H3K9me2 levels for the *c-myc* NLS construct compared to an on-slide control. **I**. 3xFLAG::MET-2 line scans in pre-gastrula embryos (1-4 cell stage) with (red) or without (grey) the *c-myc* NLS. Average of line scans across multiple nuclei are shown and error bars denote standard error of the mean. **J**. H3K9me2 levels normalized to H3 for the NLS construct (red) compared to wild-type control (grey).

Given the dependence of H3K9me2 on MET-2, we analyzed if expression of MET-2 tracked with the onset of H3K9me2. We found that MET-2 protein localization shifted from the cytosol to the nucleus from the 2-cell stage to the onset of gastrulation, and this change was observed for both endogenous MET-2 and single-copy MET-2 reporters (**Figure 2C,D, S2**). In 1-4-cell embryos, MET-2 was distributed throughout nuclei and cytoplasm with little nuclear accumulation. As embryos progressed towards gastrulation, the concentration of MET-2 within nuclei increased approximately five-fold (**Fig. 2D**). The absolute level of MET-2 protein did not change significantly over time, (**Fig. 2E**). We note that early embryos acquired nuclear MET-2 and H3K9me2 transiently during prophase, but we focus on interphase nuclei here.

Given that *met-2* is necessary for heterochromatin domains, and H3K9me2 accumulated dynamically, we asked whether regulation of MET-2 constituted part of the embryonic timer for heterochromatin establishment. We hypothesized that if nuclear MET-2 was rate-limiting, then premature accumulation of MET-2 in nuclei would lead to precocious H3K9me2 and initiate heterochromatin. We added a nuclear localization signal (NLS) from c-Myc to a FLAG-tagged copy of endogenous MET-2 using Crispr, which increased nuclear MET-2 by approximately two-fold at each stage (**Fig. 2F, I**). Increased nuclear MET-2 led to precocious accumulation of H3K9me2, beginning at least one cell division earlier than wild-type embryos (**Figure 2G, H, J**). We note that positive regulators of MET-2 may act as additional rate-limiting factors once MET-2 is nuclear, but these results indicate that gradual accumulation of MET-2 within nuclei initiates H3K9me2.

To understand how MET-2 is regulated, we searched for binding partners using immunoprecipitation followed by Multidimensional Protein Identification Technology Mass Spectrometry, using MET-2::GFP and 3xFLAG::MET-2. Wild-type *C. elegans* bearing no tagged proteins, and strains bearing an unrelated GFP or FLAG reporter served as negative controls. We chose a candidate list of interacting partners based on the specificity of MET-2 binding, on the peptide counts and protein coverage. Seventeen candidates were chosen for secondary screening for their effects on H3K9me. From this survey, two proteins emerged as likely MET-2 partners (**fig. S3A**): LIN-65 is a 100kDa protein and the most abundant interactor of MET-2 (**fig. S3B**); B0336.5 codes for a smaller protein (~30kDa) and had lower spectral counts, but similar protein coverage as LIN-65 (**fig. S3A**). We renamed B0336.5 as *arle-14* for ARL14 Effector Protein, as explained below.

To test the role of MET-2 binding partners in H3K9me deposition, we analyzed loss-of-function mutants. *lin-65* mutants had reduced levels of H3K9me1 and H3K9me2, and low, dispersed H3K9me3, similar to *met-2* mutants (**Fig. 3A-B**). *arle-14* mutants resembled a partial loss of *met-2* activity, with reduced H3K9me1/me2 levels and largely normal H3K9me3 (**Fig. 3A-B**).

**Fig. 3.**
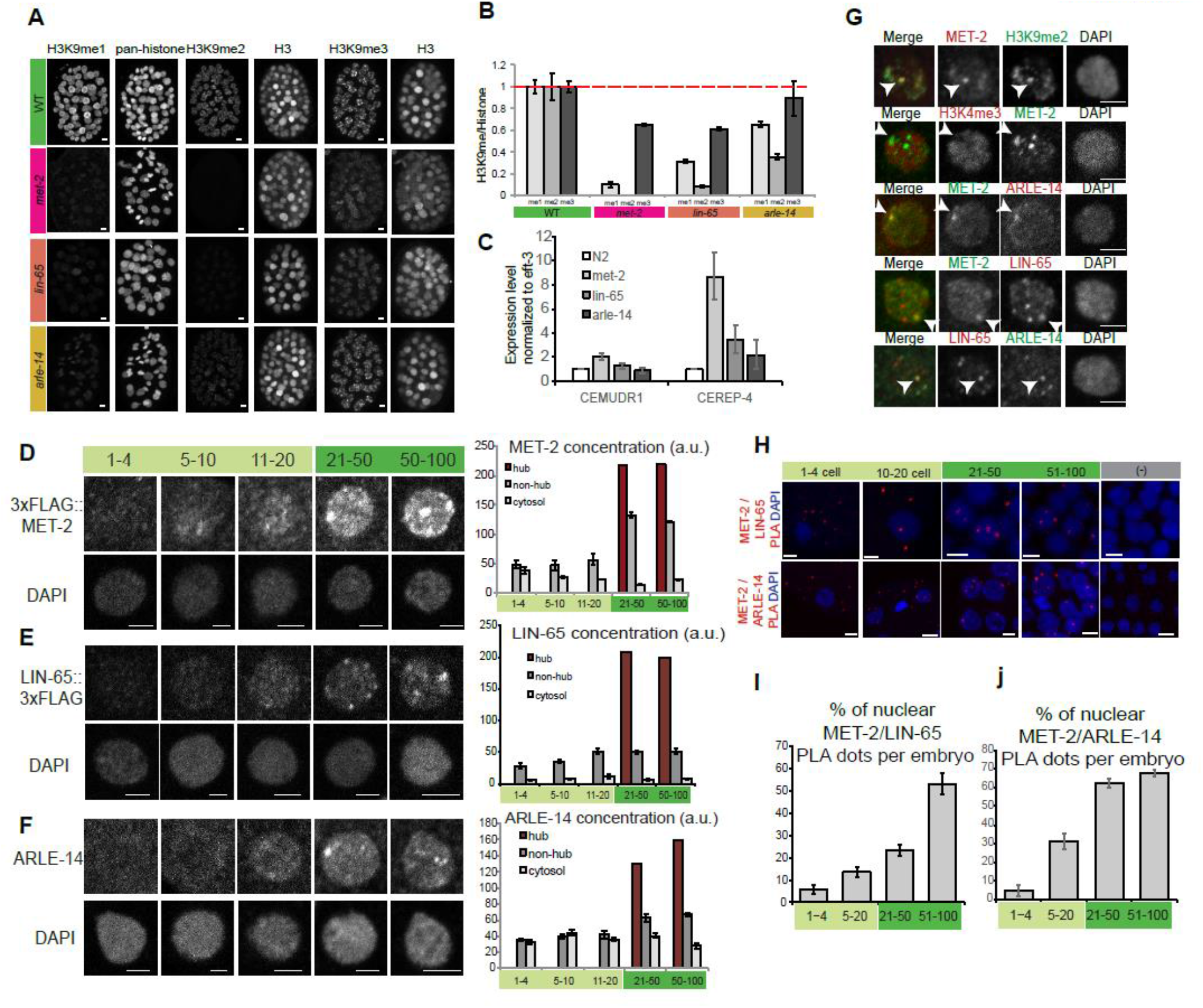
MET-2 and two conserved binding partners form concentrated nuclear hubs at gastrulation. **A**. Whole embryos stained with antibodies against methylated H3K9 and pan-histones in wild-type vs. *met-2, lin-65* or *arle-14* mutants. Scale bar, 2 μm. **B**. H3K9me levels in mutant embryos normalized to an on-slide wid-type control and to pan-histone. **C**. Expression of CEREP4 and CEMUDR1 repeat RNAs by RT-qPCR in wild-type vs. *met-2*, *lin-65* or *arle-14* mutant embryos. **D-F**. Representative singe nuclei showing 3xFLAG::MET-2 (D), LIN-65::3xFLAG (E) and ARLE-14 (F) localization at different embryonic stages (Scale bar, 2 μm). Quantitation of signal intensity in nuclear hubs (red), nuclear regions excluding hubs (“non-hub”, dark grey) and cytosol (light grey). **G**. Single nuclei showing 3xFLAG::MET-2, LIN-65::3xFLAG and ARLE-14 co-localization in concentrated protein hubs and exclusion of activating mark H3K4me3 (Scalebar 2 μm).H. PLA signal showing interactions between MET-2/LIN-65 and MET-2/ARLE-14. I-J. Percentage of nuclear PLA dots per embryos for MET-2/LIN-65 and MET-2/ARLE-14 interactions.

*Lin-65* and *arle-14* resembled *met-2* mutants in two additional assays. First, an important role of MET-2 is to silence repetitive DNA *(12, 13)*. RNAs for two repeats were de-repressed by *arle-14* mutations and, to a greater degree, by *lin-65* (**Fig. 3C**). Second, de-repressed repeats lead to a mortal germline phenotype for *met-2* mutants at 26°C *(12)*. Similarly, *lin-65* mutants became sterile after a single generation, and *arle-14* mutants after two generations. These results support the notion that LIN-65 and ARLE-14 are *bona fide* binding partners for MET-2 and contribute to its functions.

To begin to address whether LIN-65 and ARLE-14 contribute to the onset of heterochromatin formation, we examined their expression during embryogenesis. We found that both proteins behaved similarly to MET-2: both were enriched in the cytoplasm from the one-cell stage through the eight-cell stage, but gradually moved into nuclei thereafter; total levels did not change (**fig. S3C, D**). During gastrulation, we observed concentrated hubs of MET-2, LIN-65 and ARLE-14 emerge within nuclei (**Fig. 3D, E, F**). H3K9me2, MET-2, LIN-65 and ARLE-14 co-localized in many of these hubs and excluded the activating mark H3K4me3 (**Fig. 3G**).

The antibody stains suggested that MET-2 could bind LIN-65 and ARLE-14 in either the cytoplasm or the nucleus, but imaging of proteins under the light microscope lacks the resolution to define where binding occurs. We took advantage of the Proximity Ligation Assay (PLA), which detects pairs of proteins when they are within ~30 nm of each other, to investigate where MET-2 binds its partners. Positive and negative controls demonstrated that our PLA signals were specific (**fig. S3E**). At the earliest stages of embryogenesis, we observed MET-2 PLA+ signal with LIN-65 and ARLE-14 in the cytoplasm but rarely in the nucleus (**Fig. 3H, I, J**). As embryos matured, we continued to detect PLA signals in the cytoplasm but also observed signal within nuclei for MET-2 with both LIN-65 and ARLE-14. These results reveal that MET-2 interacts closely with LIN-65 and ARLE-14 and all three proteins accumulate in nuclei and nuclear hubs over time.

To address how ARLE-14 contributes to the deposition of H3K9me2 by MET-2, we examined MET-2 in wild-type and *arle-14* embryos. Neither the localization nor level of MET-2 changed in *arle-14* mutants (**fig. S4A**). By ChIP-seq, the distribution of H3K9me2 was also normal in *arle-14* mutants (wild-type vs. *arle-14* gave a genome-wide correlation of 0.8 and 0.92 for two independent experiments) (**Fig. 4A**). Thus, ARLE-14 is not required to target MET-2 in the genome (**Fig. 4B**). Rather, *arle-14* affected the degree of association of MET-2 with chromatin. Quantitative analysis by CHIP-qPCR revealed a two-fold decrease in MET-2::GFP binding to known genomic targets in *arle-14* mutants (**Fig. 4C**). Loss of *arle-14* delayed the accumulation of H3K9me2 in early embryos (**Fig. 4D, E**), revealing that ARLE-14 is also critical for proper timing of H3K9me accumulation.

**Fig. 4.**
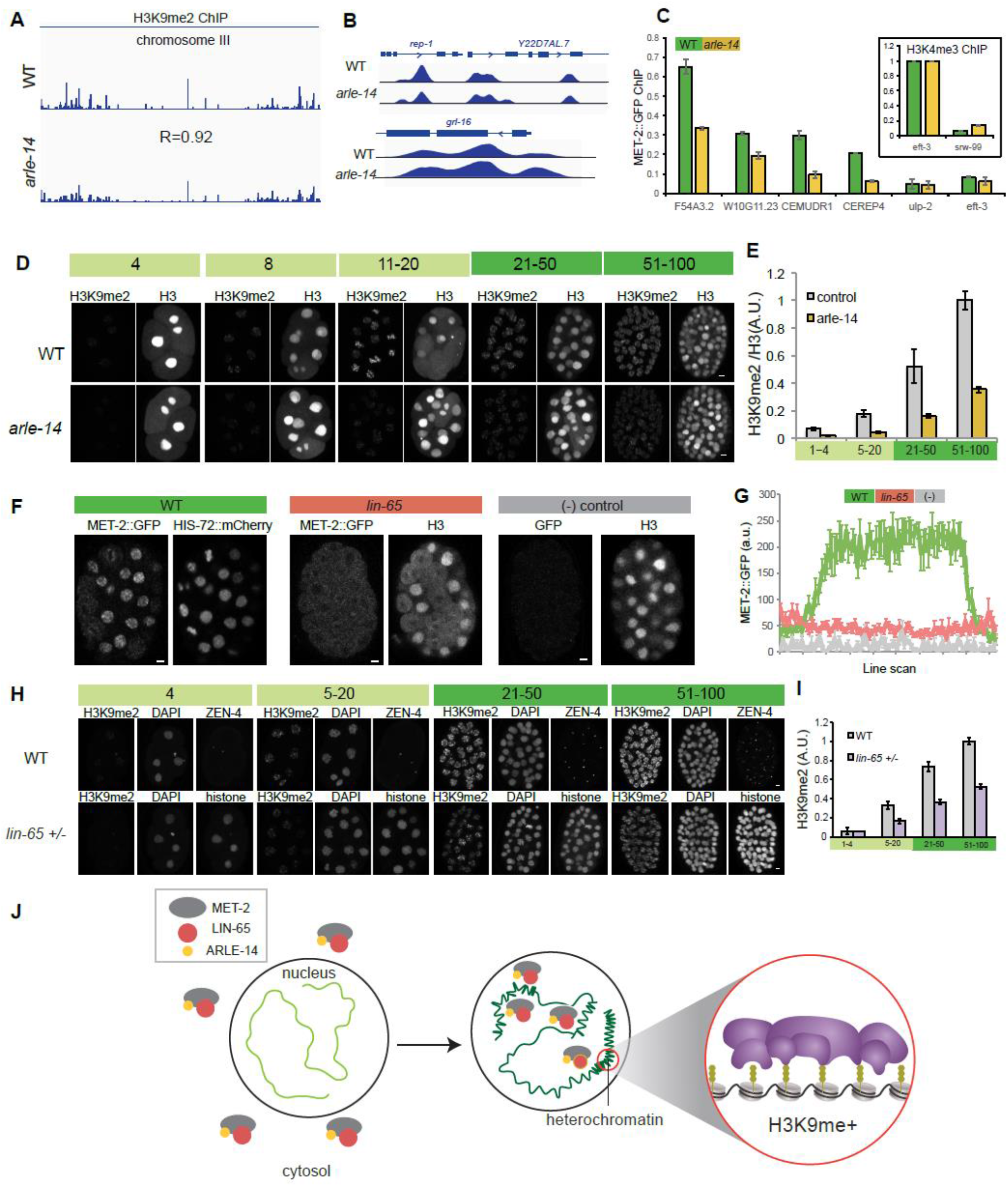
LIN-65 and ARLE-14 promote MET-2 nuclear localization and chromatin association. **A-B**. H3K9me2 ChIP-Seq (logLR) track in wild-type vs. *arle-14* mutant embryos (chromosome III and loci: *rep-1, Y22D7AL.7, grl-16).* **C**. MET-2::GFP ChIP-qPCR for wild-type (green) vs. *arle-14* (yellow) mutant embryos. Inset shows H3K4me3 ChIP-qPCR as a control. **D-E**. Acquisition of H3K9me2 in wild-type vs. *arle-14* mutants during embryogenesis, with an H3 co-stain (D) and quantitation normalized to H3 (E). **F**. Distribution of MET-2::GFP in wild-type vs. *lin-65* mutants vs. no-GFP wild-type strain (Scale bar, 2 μm). Note that H3 antibody staining is detected mostly in the cytosol during mitosis. **G**. Line-scan analysis across embryonic nuclei shows mean MET-2::GFP intensity in wild-type (green) vs. *lin-65* (pink) mutants vs. no-GFP control (“-”, grey). Average of line scans across multiple nuclei are shown and error bars denote standard error of the mean. **H**. H3K9me2 levels in embryos with a single copy ZEN-4::GFP (“WT”, on-slide control) or progeny *of lin-65 +/-* heterozygous mothers identified by histone::mCherry at designated stages of embryogenesis. **I**. Quantitation of H3K9me2 levels from wild-type (grey) or *lin-65/+* (purple) offspring. **J**. In early embryos, MET-2 (grey), LIN-65 (red) and ARLE-14 (yellow) are enriched in the cytosol, and there is little H3K9me2 or heterochromatin (light green). As embryos mature, MET-2 and interactors gradually accumulate in the nucleus, form concentrated nuclear hubs and deposit H3K9me2. MET-2-dependent H3K9 methylation is required to generate heterochromatin domains (compacted, dark green).

Next, we examined the role of LIN-65. In *lin-65* mutants, MET-2 remained cytoplasmic (**Fig. 4F, S4B**). MET-2 levels did not decrease in *lin-65* mutants (**fig. S4C**), indicating that LIN-65 affected the subcellular distribution of MET-2 and not its stability. Our results do not exclude a role for LIN-65 in regulating nuclear MET-2 but reveal that LIN-65 is required for the timely accumulation of MET-2 within nuclei.

The NLS experiment suggested that nuclear accumulation of MET-2 might be rate-limiting for H3K9me2 and heterochromatin formation. To test this idea further, we reduced the dose of MET-2 or LIN-65 by examining embryos from *met-2/+* or *lin-65/+* heterozygotes. Embryos from *met-2/+* mothers behaved like wild-type, with normal levels and distribution of MET-2 and H3K9me2 (**fig. S4D, E**), suggesting that MET-2 is dosage compensated. A half dose of *lin-65* lead to reduced accumulation of MET-2::GFP within nuclei and more cytoplasmic MET-2 (**fig. S4F**). H3K9me2 accumulated more slowly, and pre-gastrula embryos had approximately half as much compared to the wild type (**Fig. 4H-I**). These results indicate that LIN-65 is rate-limiting for nuclear accumulation of MET-2 and H3K9me.

This study makes two contributions towards understanding heterochromatin formation during embryogenesis (**Fig. 4J**). First, we have identified two binding partners, LIN-65 and ARLE-14, that are critical for MET-2 to localize to nuclei and associate with chromatin. MET-2, LIN-65 and ARLE-14 likely act together since MET-2 requires LIN-65 for nuclear localization, whereas ARLE-14 requires MET-2 for both nuclear localization and cellular accumulation (**fig S4L**).

*Lin-65* belongs to the synMuv B subclass of regulators, which are involved in chromatin regulation and transcriptional repression *(14)*. Although *lin-65* had been annotated as a novel protein *(15)*, we found similarities between LIN-65 and ATF7IP (Activating Transcription Factor 7-Interacting Protein). LIN-65 has a high-probability coiled-coil region predicted by PCOILS (CC; **fig. S4G**) and a high-confidence beta-sandwich in the C-terminus, like ATF7IP (βS; **fig. S4H**). **fig. S4I**). Like LIN-65, ATF7IP binds and localizes SETDB1 *(16–18)*. However, LIN-65 is not an obvious orthologue of ATF7IP and may be an example of convergent evolution.

Prior to this study, ARLE-14 was an uncharacterized *C. elegans* protein with homology to ADP Ribosylation Factor 14 Effector Protein (**fig. S4J**). In vertebrates, the only published function for ARL14EP is in the cytoplasm *(19).* However, the Protein Atlas shows ARL14EP in the nuclei of many human tissues (www.proteinatlas.org). We surveyed large-scale interaction databases *(20–22)* and uncovered an interaction between ARL14EP and SETDB proteins in both humans and *Drosophila* (**fig. S4K**). These data suggest that, in addition to its cytoplasmic function, ARLE-14 and its orthologues share a conserved function in nuclei with SETDB methyltransferases. Interestingly, human ARL14EP and H3K9 methylation have been implicated in polycystic ovary syndrome, but were considered independent aspects of the disease *(23–25)*. Our results suggest an intriguing link between ARL14EP and H3K9 methylation in PCOS.

A second contribution, we find that the onset of heterochromatin formation depends on the gradual accumulation of MET-2 within nuclei. Similar to *C. elegans*, mammals and *Drosophila* rebuild heterochromatin domains during embryogenesis *(1, 26)*. More generally, lack of heterochromatin domains appears to be a feature of undifferentiated cells, including embryonic stem cells and planarian neuroblasts, and differentiation involves re-establishing heterochromatin *(1)*. Examination of previous studies suggest murine SetDB1 is cytoplasmically enriched in early embryos, but its function has been difficult to address due to early lethality *(27, 28).* An intriguing idea is that localization of SETDB controls the timing of heterochromatin formation in these other animals as well.

## Acknowledgements

We thank A. Schier, D. Moazed, the Mango lab for comments on the manuscript, and S. Von Stetina for sharing the *zen-4::gfp* strain. We acknowledge support from the NIH (R37 GM056264 to SEM and R24 0D010943 to DHH), the John D. and Catherine T. MacArthur Foundation and Harvard University to SEM, the AAUW International Fellowship to BM. Some strains were provided by the CGC, funded by NIH P40 0D010440.

## Author contributions

BM and SEM designed and conducted the study and wrote the manuscript. HMC did TEM image processing. DHH did TEM. JJM, BDO and JRY performed proteomics analysis. BY, SKR and EMM generated MET-2 antibody and *met-2* Crispr and MosSCI strains. JG analyzed the H3K9me2 ChIP-Seq data. Authors have no competing interests.

## Supplementary Materials for

**Fig S1.**
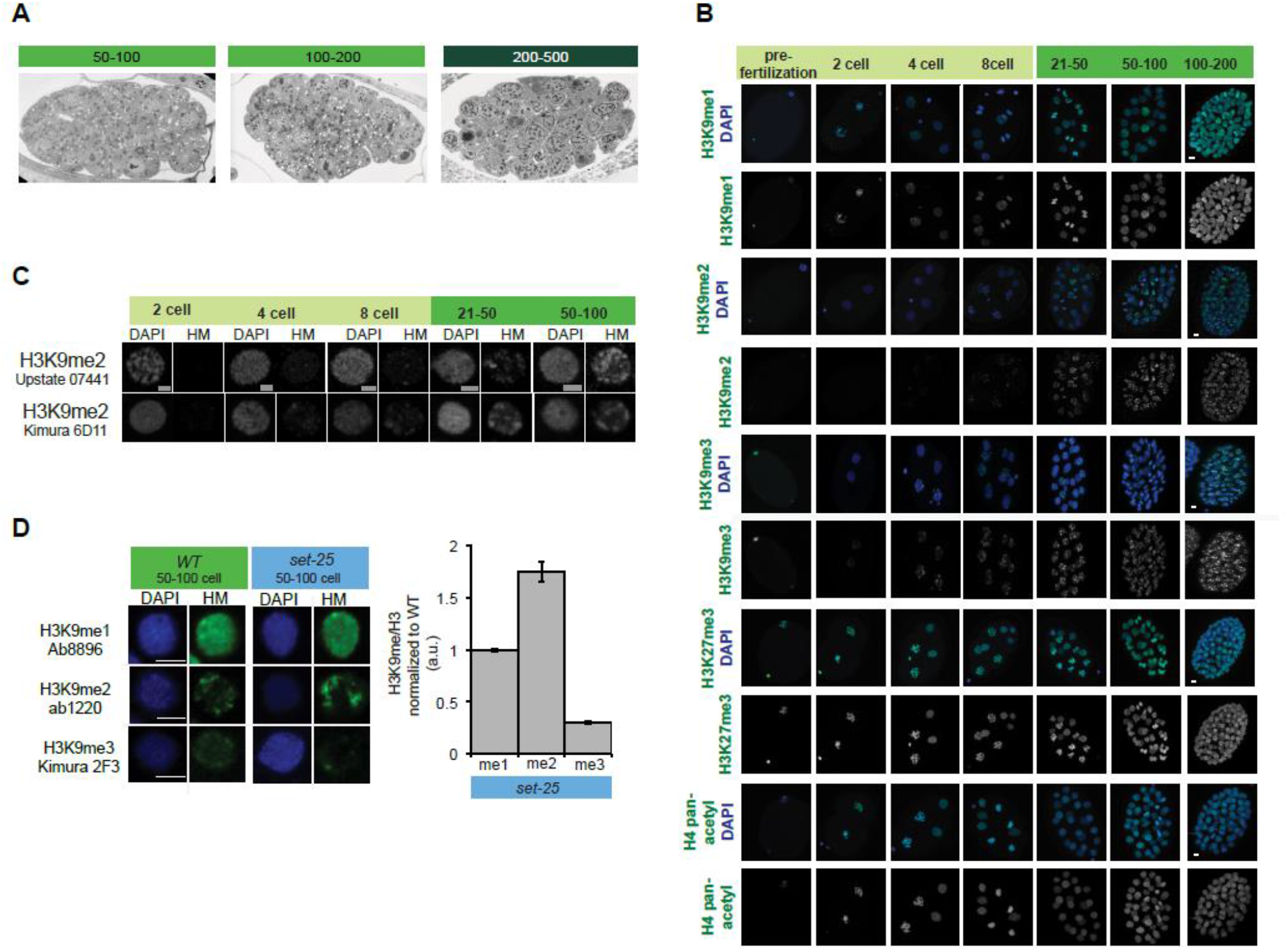
**A**. TEM single sections of whole embryos at designated stages. Each embryo is approximately 50μm long. **B**. Whole embryos stained with H3K9me1/2/3, HK27me3 or H4 pan-acetylation at different stages of development (Scale bar 2 μm). **C**. Antibody specificity for H3K9me2: representative single nuclei at designated stages showing H3K9me2 staining with two additional H3K9me2 antibodies: Upstate 07-441 and Kimura 6D11. **D**. Antibody specificity for H3K9me3: representative single nuclei at designated stages showing H3K9me staining in wild-type vs. *set-25* mutant embryos, Histone Modification (HM, green), DAPI (blue). H3K9me/Histone levels normalized to wild-type.

**Fig S2.**
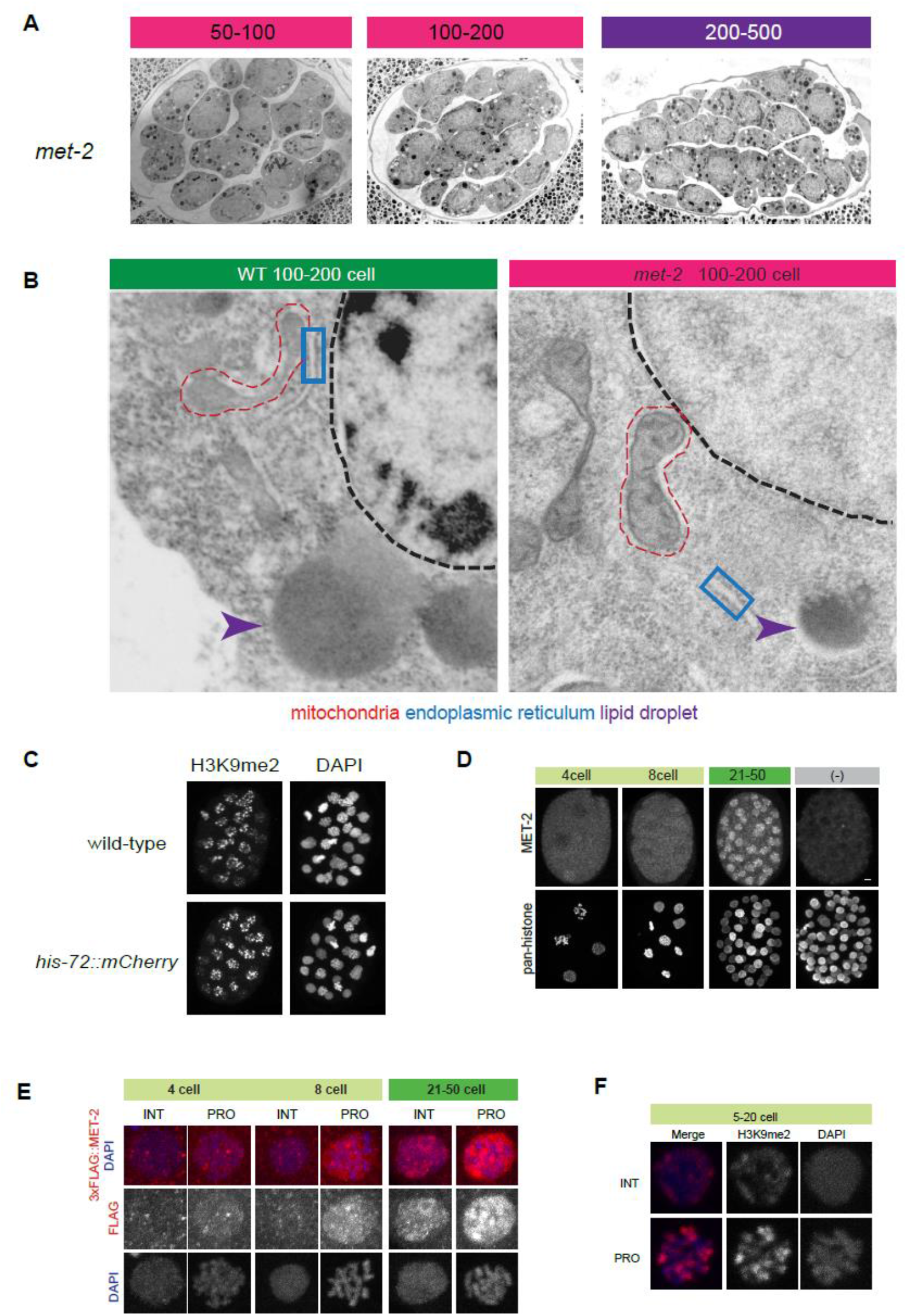
**A**. TEM single sections of whole *met-2* embryos. Note the electron dense droplets in the cytoplasm vs. the electron lucent nuclei. **B**. Cytosolic components in WT vs. *met-2* mutants by TEM. Mitochondria (red circle), endoplasmic reticulum (blue box), lipid droplet (purple arrowhead). Note the similar appearance of the cytosol vs. the different morphologies of the nuclei for wild-type vs *met-2.* **C**. H3K9me2 staining in wild-type vs. *his-72::mCherry* embryos showing that the mCherry tag doesn’t interfere with H3K9me2. **D**. Whole embryos stained with an antibody against endogenous MET-2 (raised against the first 17 amino acids of MET-2 protein) and pan-histone. (-) represents *met-2* mutants. Scale bar, 2 μm. **E**. Representative single nuclei showing Crispr reporter 3xFLAG::MET-2 during interphase (INT) and prophase (PRO, Scale bar, 2 μm). **F**. H3K9me2 levels in interphase (INT) and prophase (PRO) nuclei from the same embryo at the 15 cell stage.

**Fig S3.**
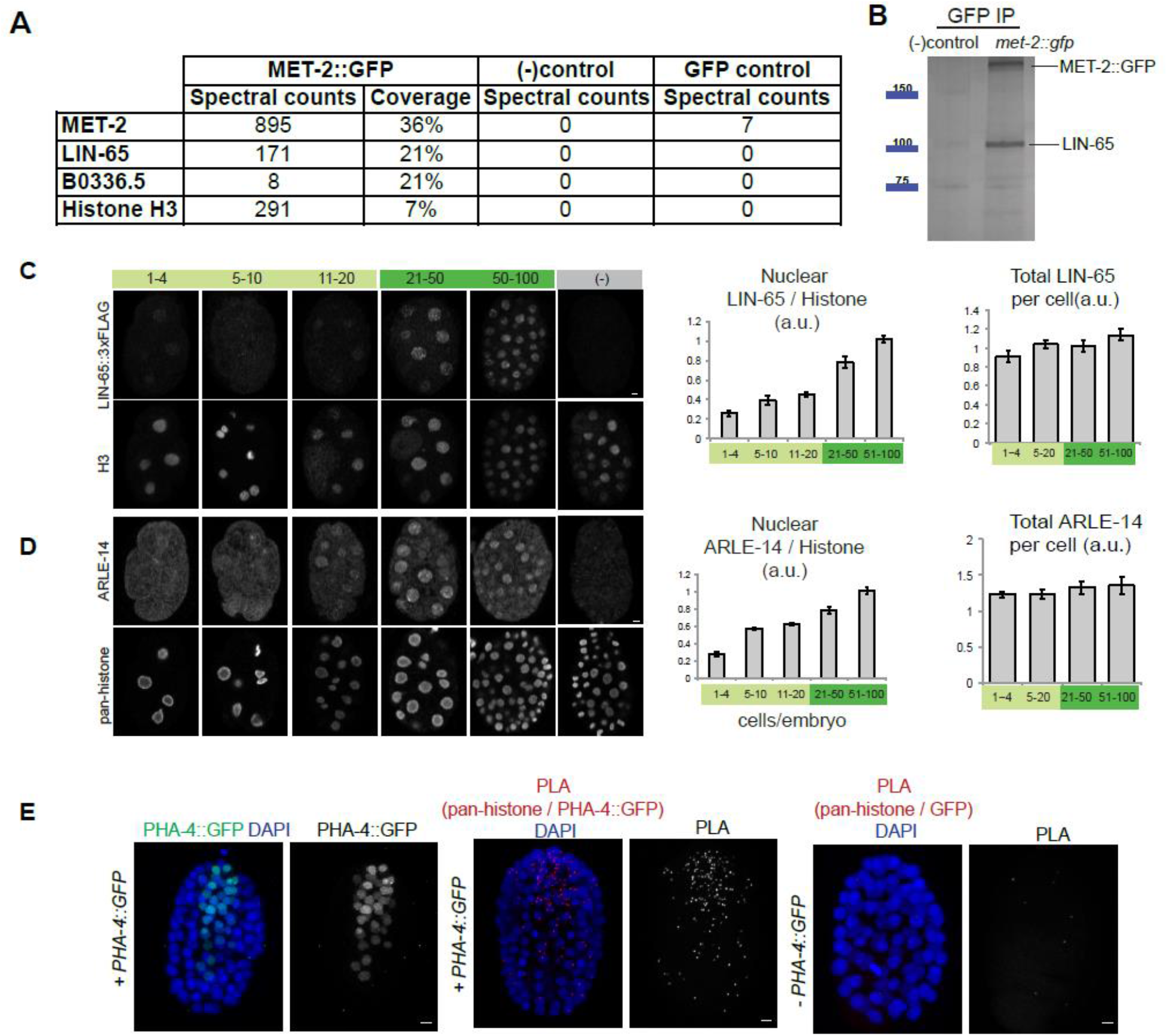
**A**. Table showing spectral counts and peptide coverage after GFP IP-MudPIT MS for the following strains: *met-2::gfp*, wild-type (no GFP) and *zen-4::gfp* (GFP control). **B**. Silver stain showing the 100 kDa LIN-65 band after MET-2::GFP IP, identified by cutting out the 100kDa band and Mass Spec. **C-D**. Whole embryos showing LIN-65::3xFLAG (αFLAG antibody; C) and endogenous ARLE-14 (D) at different stages of embryonic development with a H3 or pan-histone co-stain (Scale bar, 2 μm). Quantitation of nuclear and total protein during embryogenesis, normalized to histone. **E**. Left panel shows the staining pattern of transcription factor PHA-4::GFP (green) in whole embryos, with DNA (DAPI). The two panels on the right show PLA signal between GFP and pan-histone (red) with DNA (DAPI) in the following strains: *pha-4::gfp* or wild-type N2 *(no GFP -PHA-4::GFP)* (Scalebar 2 μm).

**Fig S4.**
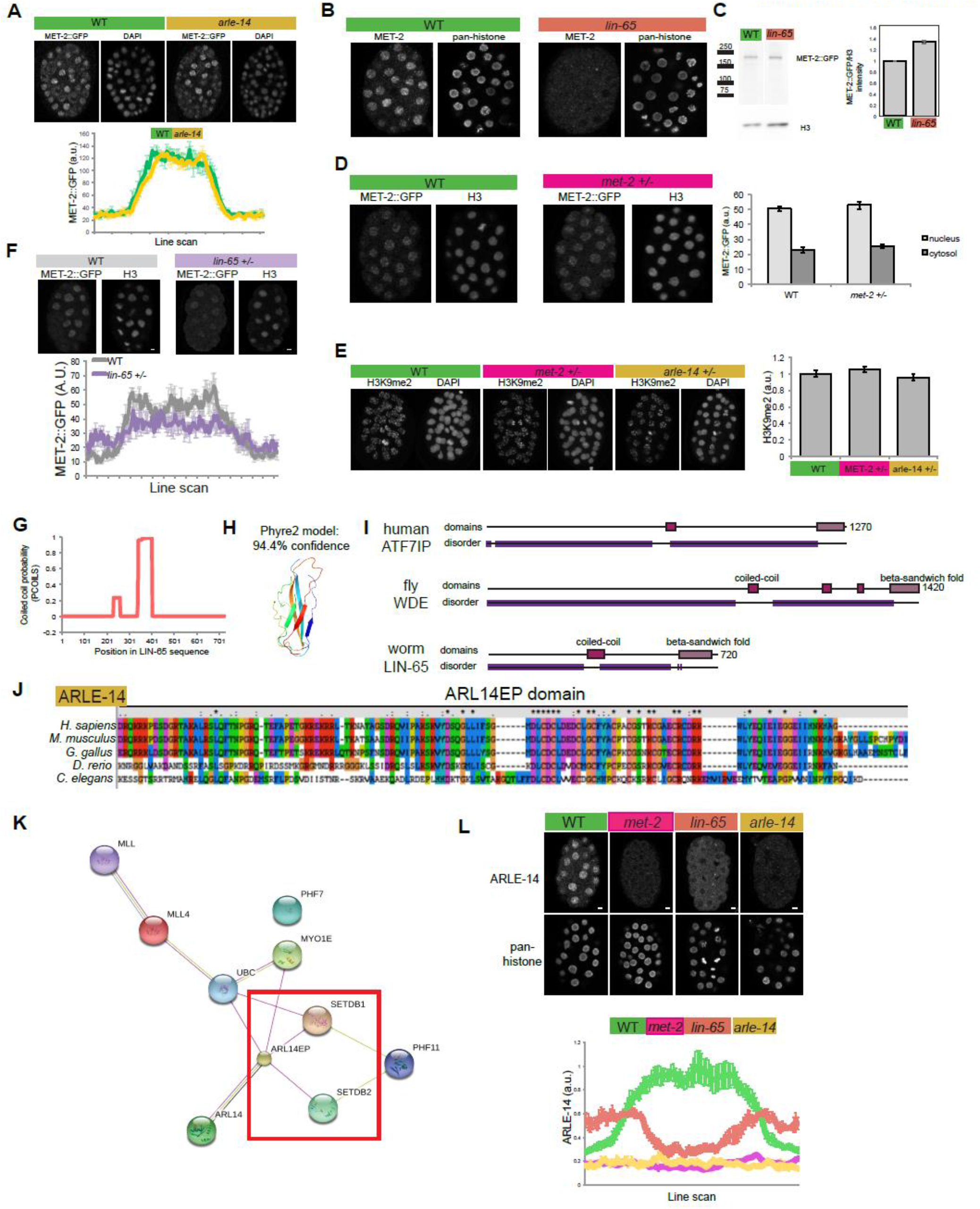
**A**. Whole embryos showing the distribution of MET-2::GFP in WT vs. *arle-14* mutants (Scale bar 2 μm). Line scan analysis showing the mean MET-2::GFP intensity across embryonic cells in WT vs. *arle-14* mutant embryos. Average of line scans across multiple nuclei are shown and error bars denote standard error of the mean. **B**. Whole embryos stained with antibodies against endogenous MET-2 protein in wild-type (green) vs. *lin-65* (red) mutants, and co-stained for pan-histone. **C**. MET-2::GFP IP and Western blot with GFP and histone antibodies in wild-type vs. *lin-65* embryonic extracts on the same gel. The lane between wild-type vs. *lin-65* is not shown. The graph shows pixel counts for the intensity of MET-2::GFP normalized to histone H3. **D**. MET-2::GFP levels in progeny of *met-2(+/-)* moms. Quantitation of MET-2::GFP signal intensity in the cytosol and the nucleus. **E**. H3K9me2 levels in progeny of *met-2(+/-)* and *arle-14(+/-)* moms and quantified. **F**. MET-2::GFP levels in WT vs. progeny of *lin-65(+/-)* heterozygous moms. Line scan quantitation showing mean accumulation of MET-2::GFP in wild-type (grey) or *lin-65/+* (purple) embryos at the 21-50 cell stage. Average of line scans across multiple nuclei are shown and error bars denote standard error of the mean. **G**. Coiled-coil probability across LIN-65 amino acid sequence predicted by PCOILS (https://toolkit.tuebingen.mpg.de/#/tools/pcoils). **H**. Model structure of LIN-65 C-terminus predicted by Phyre2 (http://www.sbg.bio.ic.ac.uk/phyre2/html/page.cgi?id=index). Full LIN-65 a.a. sequence was used as input for Phyre2. **I**. Domain architecture from HMMR (https://www.ebi.ac.uk/Tools/hmmer/) for human ATF7IP, fly Windei and worm LIN-65 showing disordered (purple), coiled-coil (pink), and beta-sandwich fold (violet) regions. **J**. Amino acid sequence alignment of worm ARLE-14 to the human ARL14EP domain using ClustalX2. Color scheme: Hydrophobic-blue, Positive charge-red, Negative charge-magenta, Polar-green, Cysteines-pink, Glycines-orange, Prolines-Yellow, Aromatic-cyan. **K**. Protein interaction map for human ARL14EP from String database (https://string-db.org/). Red box highlights interactions with human SETDB1/2. **L**. ARLE-14 antibody staining in wild-type vs *met-2, lin-65* and *arle-14* mutants. Line scan quantitation. Average of line scans across multiple nuclei are shown and error bars denote standard error of the mean.

## MATERIALS AND METHODS

**Strains**. Strains were maintained at 20°C according to *(29)*, unless stated otherwise.

*N2* (wild-type Bristol)

RB1789 *met-2 (ok2307)* III, provided by the *C. elegans* Gene Knockout Project at OMRF.

MT13232 *lin-65(n3441)* I *(15)*.

SM2078 *stIs10389 (pha-4::gfp::3xflag); pha-4 (q500) rol-9 (sc148) (30).*

SM2333 *pxSi01 (zen-4::gfp, unc-119+) II; unc-119(ed3) III.*

SM2529 *arle-14/B0336.5(tm6845) III*, provided by the Japanese National Biosource Project.

JAC500 *his-72(csb43[his-72::mCherry]) III*, provided by John Calarco *(31).*

EL597 *omIs 1 [Cb-unc-119 (+) met-2::gfp II].*

SM2491 *omIs 1 [Cb-unc-119 (+) met-2::gfp II]; met-2(ok2307) unc-119 (ed3) III.*

SM2533 *omIs1 [Cb-unc-119 (+) met-2::gfp II]; arle-14(tm6845) III.*

SM2536 *omIs1 [Cb-unc-119 (+) met-2::gfp II]; lin-65 (n3441) I.*

SM2532 *[Cb-unc-119 (+) met-2::gfp II]; his-72(csb43[his-72::mCherry]) III.*

SM2575 *lin-65::3xflag I.* This study EL634 *3xflag::met-2 III.* This study

SM2576 *arle-14(tm6845) III, lin-65::3xflag I.* This study.

SM2580 *NLS::3xflag::met-2 III.* This study.

### Number of experiments and embryos surveyed

For imaging experiments, first, many embryos were surveyed under the microscope through the eye piece and general trends noted. Then a random subset of embryos was imaged and analyzed more deeply, with quantitation. Details of analysis are described separately in the Image analysis section, and the analysis gave the same qualitative result as the trends observed in the initial survey. Below are the numbers for the quantitation. Fig.1C. >50 embryos were surveyed for each histone mark. At least 5 wild-type embryos from each developmental stage were imaged and analyzed in N=3 experiments.

**Fig. 2C**. >100 embryos were surveyed. A total of 37 embryos were imaged and analyzed in N=3 experiments.

**Fig. 2F**. >50 embryos were surveyed for each strain.14 wild-type and 14 NLS embryos were imaged and analyzed in N=3 experiments.

**Fig. 2G**. >30 more embryos for each strain were surveyed. 15 wild-type and 24 NLS embryos were imaged and analyzed in N=3 experiments.

**Fig. 3A**. We analyzed the following number of embryos in N=3 experiments: H3K9me1: WT vs. *met-2(ok2307)* (5, 7), WT vs. *lin-65(n3441)* (8, 12), WT vs. *arle-14(tm6845)* (6, 7). H3K9me2: WT vs. *met-2(ok2307)* (11, 20), WT vs. *lin-65(n3441)* (10, 13), WT vs. *arle-14(tm6845)* (22, 21). H3K9me3: WT vs. *met-2(ok2307)* (14, 12), WT vs. *lin-65(n3441)* (6, 6), WT vs. *arle-14(tm6845)* (5, 6).

**Fig. 3C**. Error bars denote standard error of the mean for N=3 experiments.

**Fig. 3H**. For MET-2/ARLE-14 PLA, a total of 20 embryos were analyzed in N=3 experiments. For MET-2/LIN-65 PLA, a total of 28 embryos were analyzed in N=3 experiments.

**Fig. 4A**. 2 biological replicates were processed in parallel.

**Fig. 4C**. Error bars denote standard error of the mean for N=3 experiments.

**Fig. 4D**. >40 embryos were surveyed for each strain.10 wild-type and 19 *arle-14(tm6845)* mutant embryos were imaged and analyzed in N=3 experiments.

**Fig. 4F**. >100 *lin-65* mutant embryos were surveyed. 14 wild-type and 31 *lin-65* mutant embryos were imaged and analyzed in N=3 experiments.

**Fig. 4H**. 22 wild-type and 25 *lin-65 +/-* embryos were analyzed in N=3 experiments.

**fig. S3C**. 41 embryos were analyzed in N=3 experiments.

**fig. S3D.** 30 embryos were analyzed in N=3 experiments.

**fig. S4A**. >30 mutant embryos were surveyed. 31 wild-type embryos and 21 *arle-14(tm6845)* mutant embryos were imaged and analyzed in N=3 experiments,

**fig. S4C.** Error bars denote standard error of the mean for N=2 experiments.

**fig. S4L**. >50 embryos were surveyed for each strain.11 wild-type vs. 11 *lin-65* mutant embryos, 8 wild-type vs. 4 *met-2(ok2307)* mutant embryos were imaged and analyzed in N=3 experiments.

### Generation of MET-2, LIN-65, ARLE-14, ZEN-4 and HIS-72 reagents

**MET-2**: To generate EL634, we inserted the 3xFLAG tag (DYKDHDGDYKDHDIDYKDDDDK) at the endogenous *met-2* locus using Crispr (32, 33). The construct places the tag at the amino terminus of MET-2 and is inserted immediately after the start codon without any linker sequences. To generate EL597, we inserted a single copy of MET-2::GFP with its endogenous promoter and upstream gene *R05D3.2* at the *ttTi5605* locus on chromosome II by MosSCI (34, 35). The GFP tag was placed at the carboxyl terminus of MET-2 and was inserted immediately before the stop codon without any linker sequences. Both the Crispr construct and the MosSCI construct could rescue H3K9me2 deposition, although the Crispr allele did so better than the MosSCI allele. The MET-2 antibody was generated against the first 17 amino acids of endogenous MET-2 and affinity purified.

To generate SM2580 (NLS::3xFLAG::MET-2), the c-Myc NLS sequence (CCAGCCGCCAAGCGTGTCAAGCTCGAC) was added directly upstream of 3xFLAG::MET-2 without any linker sequence by Crispr (36). For the insertion, the 3xFlag sequence in EL634 was targeted by the following guide RNA: ATGGACTACAAAGACCATGA(CGG). The *dpy-10* locus was used as a phenotypic marker. Because it segregated independently from the *met-2* locus, non-roller non-dpy worms were isolated for further analysis by single worm PCR and genotyping. The edit was confirmed by sequencing the 200bp region around the insertion. crRNA, tracrRNA, and Cas9 protein were ordered from the IDT Alt-R genome editing system. The 97 bp repair template was synthesized and PAGE purified by IDT.

**LIN-65**: We inserted a 3xFLAG at the endogenous *lin-65* locus using Crispr (36). The sequence of the guide RNA was TCATTCGAGAGTGATGAAGG(TGG). The 3xFLAG tag is located at the C-terminus of LIN-65 and is inserted directly before the stop codon without any linker sequences. The *dpy-10* locus was used as a phenotypic marker. Because it segregated independently from the *lin-65* locus, non-roller non-dpy worms were isolated for further analysis by single worm PCR and genotyping. The edit was confirmed by sequencing the 200bp region around the insertion. crRNA, tracrRNA, and Cas9 protein were ordered from the IDT Alt-R genome editing system. The 136 bp repair template was synthesized and PAGE purified by IDT.

**ARLE-14**: An antibody against endogenous ARLE-14 was generated. Bacteria containing *arle-14* cDNA in a pET-47b(+) (Novagen, #71461) plasmid backbone were grown at 30°C for 19 hours in LB + 50μg/ml Kanamycin. The culture was diluted 1:5 in LB + Kan and protein expression was induced with 0.25mM IPTG at 30°C for 3 hours. Bacteria were pelleted, and flash frozen at −80°C. The pellet was resuspended in 20ml Lysis Buffer (50mM Tris pH 7.2, 300mM NaCl, 5mM B-ME, 10% glycerol) and digested with 200μl of lysozyme (50mg/ml, Thermo Fisher Scientific #90082) for 30 minutes on ice. Following lysozyme digestion and sonication on ice (4 cycles, 30sec ON, 1min OFF. Output Control 3, Duty Cycle 50%, Pulsed), the protein was purified from inclusion bodies as follows: The sample was centrifuged at 4000rpm at 4°C for 15min and the supernatant discarded. The pellet was resuspended in 20ml Lysis Buffer with 1% Triton-X and 200μl Turbo DNAse (Thermo Fisher Scientific AM2239) and incubated for 20 min at room temperature. The sample was sonicated and centrifuged again with the same settings as before, and the supernatant discarded. The pellet was rinsed once with Dilution Buffer (10mM Tris/Cl pH 7.5,150mM NaCl, 0.5mM EDTA) and resuspended in Denaturation Buffer (50mM Tris-HCl pH 8, 300mM NaCl, 2mM B-ME, 5mM MgCl2, 6M urea) by gentle rocking on a shaker at room temperature for 1 hour. The solution was dialyzed against 50mM Tris-Cl pH 8, 150mM NaCl, 5mM MgCl2 and 1mg/ml protein was sent to Covance for injections into rabbits. Total IgG purification was performed by Covance after the final bleed. For *in vivo* imaging, the antibody solution was pre-cleared overnight with *arle-14(tm6845)* mutant embryos before use. Protocol described in the antibody staining section was followed to prepare *arle-14(tm6845)* embryos and stain them with the ARLE-14 antibody. The resulting pre-cleared antibody solution was transferred to a fresh tube, stored at 4°C for < 1week and used in staining experiments.

**ZEN-4**: ZEN-4::GFP was amplified from bsem1129 *(37)* with zen-4_uni_5’_nested_2_attB1 and unc-54_3’UTR_Hobert_nested_3’_attB2 primers using Takara PrimeStar (37). The resulting attB-flanked PCR product was recombined into pCFJ151 (Addgene) using Gateway BP Clonase II (Invitrogen/Thermo Fisher) to create bsem1267. SM2333 was generated by injecting bsem1267 along with pCFJ601, pMA122, pGH8, pCFJ90 and pCFJ104 (all available from Addgene) into SM2288 (ttTiS605 II; unc-119(ed3)III). The mosSCI protocol on wormbuilder.org was used to generate single integrants (34). SM2333 was used as an on-slide wild-type control in antibody stains.

**HIS-72**: The mCherry tag was inserted at the C-terminus of the endogenous *his-72* locus by Crispr *(31).* Briefly, JAC499 was injected with Cre recombinase to remove the selection cassette and produce a functional HIS-72::mCherry protein. JAC500 *his-72(csb43[his-72::mCherry]) III* was used as a histone control in MET-2::GFP stains (Fig. 2C) and to mark cross-progeny after mating (Fig. 4H, S4E). The mCherry tag did not interfere with H3K9me2 (fig. S2C).

**Antibody staining**. Antibody staining was performed as described previously (Kiefer et. al, 2007). The following antibodies were used for immunostaining with 5min 2% paraformaldehyde (PFA), 3 min methanol (for all except ARLE-14 and MET-2, which was fixed with 10min 2% PFA, 3min methanol). For mutant configurations, an on-slide wild-type sample was included, marked with a single-copy ZEN-4::GFP tag. On-slide controls allowed better quantitation between different genotypes or stages.

H3K9me1 (1:200) Abcam ab8896

H3K9me2 (1:200) Abcam ab1220, Kimura 6D11 - MABI0307

H3K9me3 (1:200) Kimura 2F3 - MABI0308

H3K27me3 (1:200) Active Motif 61017

H4-pan acetyl (1:500) Active Motif 39925

Pan-histone (1:500) Chemicon/Millipore MAB052

Histone H3 (1:500) Abcam ab1791

H5 Polymerase II (1:100) Covance MMS129-R

FLAG M2 (1:100) Sigma F1804

GFP (1:500) Millipore Sigma MAB3580

GFP (1:500) ThermoFisher Scientific A11122

MET-2 (1:500) Raised against the first 17 amino acids of MET-2 and affinity purified ARLE-14 (1:500)

**Proximity Ligation Assay**. Sigma Duolink In Situ Kit (DUO92101) was used for this assay. Embryos were fixed as for regular antibody staining. After overnight staining with primary antibodies at 15°C, the sample was stained at 37°C for 1 hour with secondary antibodies that have oligonucleotide probes attached. Connector oligos were hybridized to the probes and served as templates for circularization by enzymatic ligation when in close proximity. The ligation reaction was incubated at 37°C for 30 minutes. The circularized DNA strands were used for rolling circle amplification (RCA) and the RCA product was detected by hybridizing fluorescently labeled oligos. The RCA reaction was incubated at 37°C for 100 minutes. After each step, slides were washed with TBS + 0.2% TritonX for 5 minutes. Slides were mounted in DAPI.

### Image analysis

**Quantitation of histone modifications**. Stacks of optical sections were collected with a ZEISS LSM700 or LSM880 Confocal Microscope and analyzed using Volocity Software. Nuclei were identified in 3-D using the DAPI channel, and sum signal intensity of histone modification was calculated for each nucleus in interphase. Mitotic nuclei were excluded from the analysis manually based on DAPI morphology. Mean cytoplasmic background was measured at a random point for each embryo, and mean background was subtracted from the nuclear signal by multiplying with nuclear volume. For each nucleus, signal intensity of histone modification was normalized to signal intensity of histones. Normalized values were averaged for nuclei at given stages and plotted. (Fig. 1D, 2D, 2E, 2J, 3B, 4E, S1D, S3C, S3D).

**TEM**. Adult hermaphrodites were rapidly chilled in liquid nitrogen while exposed simultaneously to very high pressure (2100 bar). This combination preserves the morphology of organelles and finer structures. Frozen samples underwent freeze substitution to deposit osmium tetroxide for contrast and fixation. Worms were embedded and sectioned along the ovary to view multiple embryos in a row. A detailed protocol is available in (5).

**TEM Image processing**. Raw images were processed with Photoshop as 8-bit gray scale images. Images from different preparations were standardized for accurate comparison. The cytoplasm was used to adjust signal intensity range for each image, but image contrast was not altered. Adjusted images were then saved and quantified with Image J by Line scan analysis.

**TEM Line scan analysis**; Random lines were drawn across the center of different nuclei, and the intensity measured. Standard deviation was calculated for each individual line. Standard deviation of 30 lines were averaged for each strain and listed on the plot. A randomly selected line profile for a nucleus is shown as an example. The standard deviation describes the morphology of the nucleus, i.e. a higher standard deviation stems from a more punctate staining pattern that alternates between high and low values (ie electron dense heterochromatin and electron lucent nucleoplasm).

**MET-2/ARLE-14 Line scan analysis**: Lines that go through the center of the nucleus were drawn across the cell, and the intensity measured. Each line had 100 bins. The intensity in each bin was averaged for 30 lines, and the average line plotted. Error bars denote the standard error of the mean at each bin (Fig. 2I, 4G, S4A, S4F, S4L). Definition and Quantitation of hubs. Nuclear hubs were defined by intensity thresholding in ImageJ. The threshold was selected manually in wild-type nuclei at the 51 −100 cell stage and the same threshold was applied to all the images in a given dataset. The intensity measurements for each defined hub were averaged to yield the intensity of “hubs” at given embryonic stages. “Non-hub” was defined as nuclear areas that were below the intensity threshold. The mean intensity in non-hub areas was measured for each nucleus and averaged across 30 nuclei. Non-interphase nuclei were discarded manually. For intensity measurements in the cytosol, at least 4 random areas in the cytosol was chosen for every cell that was in interphase. The measured intensities were averaged.

**Half-dose LIN-65 experiments**: *lin-65 (n3441)* moms were crossed with JAC500 *his-72::mCherry* males. Progeny of *lin-65+/-* heterozygotes marked by mCherry were analyzed. On the same slide, SM2333 containing a single copy of *zen-4::gfp* was used as a wild-type staining control. Mean H3K9me2 intensity in each nucleus was quantified using Volocity and the average H3K9me2 intensity per nucleus plotted, normalized as described in the staining section.

### Biochemistry

**Harvesting embryos**. Mix-staged embryos were collected from adult worms by bleaching Embryos were, were shaken at 200rpm at 20°C in CSM medium (100mM NaCl, 5.6mM K2HPO4, 4.4mM KH2PO4, 10μg/ml cholesterol, 10mM potassium citrate, 2mM CaCl2, 2mM MgSO4, 1X trace metals.) without food (38). Once the embryos hatched and became L1s, concentrated NA22 bacteria were added to the culture. Synchronized embryos were harvested by bleaching after 62-66 hours when most worms carried 1-8 embryos, frozen in liquid nitrogen and stored at −80°C.

**Immunoprecipitation**. Frozen embryo pellets were resuspended in Lysis Buffer (50mM HEPES pH 7.4, 1mM EGTA, 1mM MgCl2, 100mM KCl, 10% glycerol, 0.05% NP40) with protease inhibitors (Calcbiochem Cocktail Set I, #539131) and incubated on ice for 10 minutes. Using the QSonica Q800 Sonicator, samples were sonicated at 40% amplitude, 10 seconds on, 50 seconds off, for 3 cycles. After sonication, samples were centrifuged for 10 minutes at 10,000g at 4°C. The supernatant was transferred to a new tube and diluted in Dilution Buffer (10mM Tris-Cl pH 7.5, 150mM NaCl, 0.5mM EDTA). 1.5 mg of total lysate was pre-cleared for 1 hour at 4°C with 25 μl Chromotek magnetic agarose beads prior to IP. The pre-cleared lysate was then used for immunoprecipitating MET-2::GFP With Chromotek GFP-TRAP magnetic agarose beads or Sigma FLAG M2 antibody coupled to magnetic agarose beads for 5 hours at 4°C. Beads were rinsed with Dilution Buffer 3 times, washed with Dilution Buffer twice for 5 minutes, and eluted with 50μl 0.2M glycine pH 2.5 for 30 seconds under constant mixing or with 200μg/ml 3xFLAG peptide in TBS for 1 hour at 4°C. 5μl 1M Tris base pH 10.4 was added for neutralization after glycine elution. Samples were boiled in 2X Laemmli Sample Buffer (Biorad #161-0737) with 50mM DTT and analyzed by Western blotting or silver staining (SilverQuest Silver Staining Kit, ThermoFisher, LC6070).

**Mass spectrometry**. *Reagents and Chemicals;* Deionized water (18.2 MΩ, Barnstead, Dubuque, IA) was used for all preparations. Buffer A consists of 5% acetonitrile 0.1% formic acid, buffer B consists of 80% acetonitrile 0.1% formic acid, and buffer C consists of 500 mM ammonium acetate and 5% acetonitrile.

***Sample Preparation***; Proteins were precipitated in 23% TCA (Sigma-Aldrich, St. Louis, MO, Product number T-0699) at 4 °C O/N. After 30 min centrifugation at 18000 × g, protein pellets were washed 2 times with 500 ul ice-cold acetone. Air-dried pellets were dissolved in 8 M urea/ 100 mM Tris pH 8.5. Proteins were reduced with 1 M Tris(2-carboxyethyl)phosphine hydrochloride (Sigma-Aldrich, St. Louis, MO, product C4706) and alkylated with 500 mM 2-Chloroacetamide (Sigma-Aldrich, St. Louis, MO, product 22790-250G-F). Proteins were digested for 18 hr at 37 °C in 2 M urea, 100 mM Tris pH 8.5, 1 mM CaCl2 with 2 ug trypsin (Promega, Madison, WI, product V5111). Digestion was stopped with formic acid, 5% final concentration. Debris was removed by centrifugation, 30 min 18000 × g.

***MudPIT Microcolumn**;* A MudPIT microcolumn *(39)* was prepared by first creating a Kasil frit at one end of an undeactivated 250 μm ID/360 μm OD capillary (Agilent Technologies, Inc., Santa Clara, CA). The Kasil frit was prepared by briefly dipping a 20 - 30 cm capillary in well-mixed 300 μL Kasil Kasil 1624 (PQ Corporation, Malvern, PA) and 100 μL formamide, curing at 100°C for 4 hrs, and cutting the frit to ~2 mm in length. Strong cation exchange particles (SCX Partisphere, 5 μm dia., 125 Å pores, Phenomenex, Torrance, CA) was packed in-house from particle slurries in methanol 2.5 cm. Additional 2.5 cm reversed phase particles (C18 Aqua, 3 μm dia., 125 Å pores, Phenomenex) were then similarly packed into the capillary using the same method as SCX loading, to create a biphasic column. An analytical RPLC column was generated by pulling a 100 μm ID/360 μm OD capillary (Polymicro Technologies, Inc, Phoenix, AZ) to 5 μm ID tip. Reversed phase particles (Aqua C18, 3 μm dia., 125 Å pores, Phenomenex, Torrance, CA) were packed directly into the pulled column at 800 psi until 12 cm long. The MudPIT microcolumn was connected to an analytical column using a zero-dead volume union (Upchurch Scientific (IDEX Health & Science), P-720-01, Oak Harbor, WA).

***LC-MS/MS analysis*** was performed using an Agilent Technologies 1200 HPLC pump and a Thermo Orbitrap Velos using an in-house built electrospray stage. MudPIT experiments were performed with steps of 0% buffer C, 30% buffer C, 50% buffer C, 90/10 % buffer C/B and 100% C, being run for 3 min at the beginning of each gradient of buffer B. Electrospray was performed directly from the analytical column by applying the ESI voltage at a tee (150 μm ID, Upchurch Scientific)(39). Electrospray directly from the LC column was done at 2.5 kV with an inlet capillary temperature of 325 °C. Data-dependent acquisition of MS/MS spectra with the Orbitrap Velos were performed with the following settings: MS/MS on the 10 most intense ions per precursor scan; 1 microscan; reject unassigned charge state and charge state 1; dynamic exclusion repeat count, 1; repeat duration, 30 second; exclusion list size 200; and exclusion duration, 30 second.

***Data Analysis**:* Protein and peptide identification and protein quantitation were done with Integrated Proteomics Pipeline - IP2 (Integrated Proteomics Applications, Inc., San Diego, CA. http://www.integratedproteomics.com/). Tandem mass spectra were extracted from raw files using RawConverter *(40)* with monoisotopic peak option and were searched against Wormbase protein database (WB257) with reversed sequences using ProLuCID (41, 42). The search space included all half and fully-tryptic peptide candidates. Carbamidomethylation (+57.02146) of cysteine was considered as a static modification. Peptide candidates were filtered using DTASelect with the parameters -p 2 -y 1 --trypstat --pfp 0.01 --extra --pI -DM 10 --DB --dm -in -m 1 -t 2 --brief --quiet (43). Mass spectrometry identified a total of 602 proteins in the MET-2::GFP sample. 500 proteins were eliminated due to their presence in controls, and 43 proteins were eliminated because they had low sequence coverage (<6%). Remaining 59 proteins were further analyzed based on suggested function and localization from literature. Promising candidates were chosen for a loss-of-function screen.

**RNA expression analysis**. Embryos frozen in liquid nitrogen were partially thawed. An equal volume of glass beads (Sigma G8772) were added and samples were vortexed for a minute. For Trizol-chloroform extraction, 1ml Trizol (Thermo Fischer Scientific #15596026) was added to the sample, vortexed for 30 seconds and incubated at room temperature for 5 minutes. 300 μl chloroform (Calbiochem, Omnipur #3155) was added, shaken by hand and incubated at room temperature for 3 minutes. The sample was centrifuged at 14000rpm at 4°C for 15 minutes. Upper aqueous phase was transferred to a fresh tube and RNA was precipitated using 500μl isopropanol and 1 μl Glycoblue (Thermo Fisher Scientific AM9516). The pellet was air-dried and resuspended in nuclease-free water at room temperature for 10 minutes. Samples were treated with DNAse at 37°C for 30 minutes using the Turbo DNA-free kit (Thermo Fisher Scientific AM1907), and 500 ng of RNA was used for reverse transcription with a random primer mix (Protoscript First Strand cDNA Synthesis Kit, NEB E6300). The synthesis reaction was diluted in water to yield a total volume of 50μl, and 3μl of the cDNA was analyzed by qPCR (KAPA SYBR FAST qPCR Master Mix, KK4601).

**ChIP**. H3K9me2 ChIP was done as described previously (30) with the following changes: Embryos frozen in liquid nitrogen were thawed on ice and fixed in 1.5% Formaldehyde (Electron Microscopy Sciences, #15686) for 15 minutes at room temperature. Using the QSonica Q800 Sonicator, samples were sonicated at 30% amplitude, 30 seconds ON, 30 seconds OFF for a total of 15 minutes at 4°C, yielding 100-300 base pair fragments. 6μl of Kimura 6D11 antibody was coupled to 25μl beads (Pierce Protein A/G magnetic beads, #88802) for 8 hours at 4°C prior to ChIP. 40 μg of chromatin was used per ChIP reaction and chromatin was pre-cleared for 2 hours at 4°C using uncoupled magnetic beads (Pierce #88802). To elute the bound immunocomplexes, 150 μL of elution buffer (50mM NaHCO3, 140mM NaCl, 1% SDS) was added to each tube and heated at 65°C for 15 minutes. For MET-2::GFP ChIP, embryos were fixed with 1,5mM EGS (Pierce #21565) for 10 minutes and with 1% Formaldehyde for another 10 minutes. 6μl of GFP antibody (Abcam5665) was coupled to 25μl beads for 8 hours at 4°C prior to ChIP. 2μl of H3K4me3 (Abcam8580) antibody was coupled to 25μl beads for 8 hours at 4°C prior to ChIP.

***Library preparation***: The ChIP-Seq libraries were generated by using Apollo 324 System and PrepX ILM DNA Library Kit from IntergenX. After adaptor ligation, the input and ChIP DNA were enriched by PCR amplification using NEBNext High-Fidelity 2X PCR master Mix with Q5 polymerase and PrepX PCR primer with the following PCR conditions: 30 seconds at 98°C, [10 seconds at 98°C, 30 seconds at 60°C, 30 seconds at 72°C] for 8 cycles for Input libraries and 11 cycles for ChIP libraries, following 5 minutes at 72°C (~15 uL adaptor ligated DNA, 25 uL NEBNext High-Fidelity 2X PCR master Mix, 2uL Universal PCR primer, brought to to 50ul with Nuclease-Free water). The enriched DNA was then purified using 50 uL (1:1 ratio of DNA volume and beads) of PCR Clean DX Beads (Aline) and size selected by Pippin Prep, 180-600bp. 1 uL of each library was applied to measure the concentration using a Qubit dsDNA assay kit (Invitrogen). 1 ng of DNA from each library was checked by a TapeStation (Agilent Technologies). Input and ChIP libraries were pooled such that they each had the same amount of molecules and expected for obtaining the similar number of reads. The Illumina sequencing was performed with 75 nt paired-end reads.

***Sequencing Analysis***: DNA fragments were sequenced on an Illumina HiSeq machine, yielding 64-95 million 75bp paired-end reads per sample. Reads were aligned to the *C. elegans* reference genome (ce10) with Bowtie2 (44), version 2.3.1, and default parameters except for ‘--very-sensitive’ and ‘-X 2000’. PCR duplicates were removed from the alignment files after identifying properly paired fragments that shared both leftmost and rightmost genomic coordinates. MACS2 (45), version 2.1.1.20160309, was used to call peaks in the ChIP samples, using input DNA as the control and analyzing only properly paired fragments (-f BAMPE). Peaks were annotated using ChIPseeker (46), version 1.12.0. Log-likelihood ratio tracks were calculated using the MACS2 module ‘bdgcmp’, and correlations were calculated using the BigWig tools ‘bedGraphToBigWig’ and ‘wigCorrelate’ (47).

***Data access***. The data from this study will be submitted to NIH SRA and GEO database with accession numbers (pending).

